# Sensory processing sensitivity is associated with neural synchrony and functional connectivity during threatening movies

**DOI:** 10.1101/2024.03.27.586963

**Authors:** Christienne G. Damatac, Judith R. Homberg, Tessel E. Galesloot, Linda Geerligs, Corina U. Greven

**Affiliations:** Donders Institute for Brain, Cognition and Behaviour, Radboud University Medical Center, Nijmegen, The Netherlands; Department of Cognitive Neuroscience, Radboud University Medical Center, Nijmegen, The Netherlands; Donders Center for Cognitive Neuroimaging, Donders Institute for Brain, Cognition and Behaviour, Radboud University, Nijmegen, The Netherlands; Department of IQ Healthcare, Radboud University Medical Center, Nijmegen, The Netherlands; Department of Artificial Intelligence, Donders Center for Cognition, Donders Institute for Brain, Cognition and Behaviour, Radboud University, Nijmegen, The Netherlands; Karakter Child and Adolescent Psychiatry University Centre, Nijmegen, The Netherlands

**Keywords:** sensory processing sensitivity, environmental sensitivity, naturalistic, fMRI, neuroimaging

## Abstract

Sensory processing sensitivity (SPS) is an evolutionarily conserved trait describing a person’s sensitivity to subtle stimuli, their depth of processing, emotional reactivity, and susceptibility to being overwhelmed. SPS is considered a fundamental and evolutionarily conserved trait, yet its neural mechanisms remain insufficiently understood. Therefore, we investigated whether SPS relates to processing movies differently in the central executive (CEN), default mode (DMN), and salience (SN) networks. We obtained positive and negative dimension Sensory Processing Sensitivity Questionnaire (short-form) scores and (neutral and threat aural framing) movie-fMRI data from a population-based sample (Healthy Brain Study, N=238, age_mean_=34years). We performed *a priori* inter-subject representation similarity, activation, and inter-subject functional connectivity analyses to characterize SPS-dimension-related neural responses during movie-viewing. More similar negative dimension SPS score related to more neural synchrony in the CEN and SN during threat. Higher negative dimension SPS score related to reduced CEN-DMN functional connectivity during threat, an effect shared across between-network regions but most strongly driven by reduced connectivity between right dorsomedial prefrontal cortex and left lateral prefrontal cortex. Our findings suggest that highly sensitive individuals exhibit distinct CEN differences shaping environmental perception, process threat differently, and each SPSQ-SF dimension may involve unique neurological mechanisms.

## 1. INTRODUCTION

Individuals exhibit varying degrees of sensitivity to both positive and negative environmental stimuli, as described by the framework of environmental sensitivity (Pluess, 2015). One relevant trait capturing inter-individual differences in environmental sensitivity is Sensory Processing Sensitivity (SPS), defined as a person’s sensitivity to subtle stimuli, their depth of processing, emotional reactivity, empathy, and susceptibility to being overwhelmed (Aron & Aron, 1997). SPS is a fundamental, evolutionarily conserved trait (Pluess, 2015) and individual differences stem from a combination of genetic and environmental factors (Aron et al., 2012; Belsky & Pluess, 2009; Greven et al., 2019; Homberg et al., 2016). Also known as “high sensitivity,” higher SPS has been associated with a plethora of observed health-related outcomes: more internalizing problems, burnout, anxiety, and depression symptoms; more displeasure with work and need for recovery; less subjective happiness, life satisfaction, stress-management, and emotion regulation; more ill-health symptoms and nonprescription medication use; and more susceptibility to distractions, moderating memory retention (Andresen et al., 2018; Bakker & Moulding, 2012; Benham, 2006; Booth et al., 2015; Brindle et al., 2015; Damatac et al., 2023; De Gucht et al., 2022; Evers et al., 2008; Iimura & Takasugi, 2022; Jonsson et al., 2014; Liss et al., 2005, 2008; Marhenke et al., 2023; Meredith et al., 2016; Neal et al., 2002; Sobocko & Zelenski, 2015; Yano & Oishi, 2018). Therefore, characterizing the underlying neural mechanisms of SPS can be informative for improving symptom prevention and health promotion in relation to environmental sensitivity.

In recent decades, SPS has gained increasing attention in the field of psychology, leading to the development of a more comprehensive questionnaire assessment; the 43-item Sensory Processing Sensitivity Questionnaire (SPSQ) (De Gucht et al., 2022), which overcomes the limitations of the traditional questionnaire, the Highly Sensitive Person Scale (Aron & Aron, 1997). The SPSQ and its shorter form (De Gucht & Woestenburg, 2024), used here, include two dimensions with corresponding subscales: a positive (*aesthetic sensitivity, sensory comfort, sensory sensitivity to subtle internal and external stimuli,* and *social-affective sensitivity*) and negative (*emotional and physiological reactivity* and *sensory discomfort*) one. Thus, we may expect that the positive versus negative dimensions reflect distinct patterns in association with brain responses. Notably, as research on SPS is still in its infancy, the dimensionality of SPS and its neural mechanisms remain insufficiently understood.

Functional magnetic resonance imaging (fMRI) may elucidate SPS-related brain mechanisms as a method for testing sensitivity-related susceptibility to environmental influence in a controlled setting. Some fMRI evidence for its neural basis exists: SPS may characterized by a “hypersensitive” brain, reflected by more activation in areas implicated in the response to social-emotional and other environmental stimuli (Acevedo et al., 2014, 2017, 2021a; Aron et al., 2012; Chen et al., 2015). Subtle stimuli awareness and processing depth are signature features of SPS wherein highly sensitive individuals may perceive their environments differently. Their perceptual capacity may be larger so that the threshold for a stimulus to enter awareness might be reduced or associative thinking processes might be stronger (e.g., quicker stimulus detection, related to high emotional reactivity) (Acevedo et al., 2018, 2021a; Greven et al., 2019). Together, brain activation patterns in high SPS individuals indicate deep information processing, as suggested by increased activation in the precuneus, prefrontal cortex, inferior frontal gyrus. They also suggest increased emotionality and empathy, as reflected in increased activation in the insula, claustrum, amygdala, cingulate cortex during social affective tasks (Acevedo et al., 2014, 2017, 2021a; Chen et al., 2015; Jagiellowicz et al., 2011). The aforementioned clusters of brain regions correspond to the default mode (DMN) and salience (SN; ventral attention) networks which mediate internal mentation and attention towards salient and emotional stimuli, respectively (Seeley, 2019; Smallwood et al., 2021). Brain regions are not active in isolation, but jointly, and each network has different functions; SN regulates attention to important stimuli, DMN engages during introspection and self-referential processing, and the central executive network (CEN) orchestrates cognitive control and task management. Based on former fMRI studies on SPS, one may expect to observe environment-related changes in functional connectivity as a function of an individual’s degree of sensitivity. Conceptualized in a neural model by (Homberg & Jagiellowicz, 2021), differences related to high environmental sensitivity may exist in brain regions of CEN, SN, and DMN. Based on this model, here, we solely focus on these three networks *a priori*. To further bridge this theoretical framework, we observed environment-related changes in brain function in an ecologically valid paradigm.

Movie fMRI (m-fMRI) enhances ecological validity compared to typical task-based fMRI designs and provides insight into how continuous streams of sensory stimulation can differently affect highly sensitive people. Naturalistic viewing (i.e., passive viewing of a movie) is a way to represent the complexities of emotional perception in daily life in an fMRI setting. When different people watch the same movie, their brain activities tend to synchronize, which indicates that they engage in a common mode of processing (Hasson et al., 2004, 2010; Wagner et al., 2016). Inter-subject synchrony can be measured through a metric known as inter-subject correlation (ISC), which is the similarity between each pair of subjects’ blood oxygenation level dependent fMRI activation while they view the same time-locked stimulus (Hasson et al., 2004; Jääskeläinen et al., 2008). Likewise, the subjects’ SPS questionnaire scores can be represented as the similarity between each pair of participants to generate an analogous covariance structure. Brain functional activation can then be compared to SPS scores using an inter-subject representation similarity analysis (IS-RSA) (Finn et al., 2020). For the brain regions that show a difference in ISC in relation to SPS, an interesting question is whether there are systematic increases or decreases in neural activity along the SPS spectrum, which is why we additionally analyze the neural activation in these brain regions. The underlying principle is that people who exhibit more behavioral trait similarity will also display more neural response similarity.

Here, as the first study of SPS and m-fMRI, we ask in which networks do participants with higher SPS show a different mode of processing than those with lower SPS while watching different movies. While ISC characterizes correlations between the full duration blood oxygen level-dependent signal time-course of a region between subjects, complementary inter-subject functional connectivity (ISFC) analysis can reveal complex stimulus-evoked correlation pattern *between* brain regions and has the advantage of increased signal-to-noise ratio compared to within-subject measures of functional connectivity, as the signal is driven by shared signals and not by noise (Simony et al., 2016). Moreover, compared to resting-state paradigms, naturalistic neuroimaging paradigms improve the reliability of functional connectivity estimations (Wang et al., 2017). Therefore, we additionally model associations between SPS and ISFC.

The environment exerts greater influence on highly sensitive people, wherein higher SPS has been associated with greater susceptibility to both negative, trait-inducing environments (e.g., threatening life events, stress, or overwhelming sensory input), as well as positive, health-promoting environments (e.g., daily uplifts and social support) (Aron & Aron, 1997; Damatac et al., 2023; Liss et al., 2005; Slagt et al., 2018). One study showed that high SPS participants were more emotionally reactive to positive mood induction than participants with low SPS, while another demonstrated that high SPS participants who were psychophysiologically stressed were more emotionally reactive to threatening terrorism-related pictures than their low SPS counterparts (Lionetti et al., 2018; Rubaltelli et al., 2018). Highly sensitive people may thus have greater emotional reactivity to both positive and negative environmental influences. In the present study, participants watched movies that differed by topic and aural framing. These aural framings (i.e., conditions) were neutral or threatening to the participant, allowing us to assess whether SPS dimension score correlated with neutral or threatening environmental stimuli.

In this study, we combined naturalistic IS-RSA, activation, and ISFC to characterize the underlying neural mechanisms of SPS dimensionality more fully. We analyzed cross-sectional SPS questionnaire and m-fMRI data from 238 participants from the Healthy Brain Study, a large, well-characterized healthy adult population-based sample (Healthy Brain Study consortium et al., 2021). First, we aimed to investigate the association between neural ISC and SPS dimension score similarity during each movie-viewing condition, and to subsequently test whether any significant association differed in threat versus neutral conditions. To better understand these associations, we also evaluated whether network activation differences between neutral and threat conditions are associated with SPS dimension scores. Second, we aimed to assess the association between ISFC (between and within *a priori* networks) and SPS dimension score during each movie-viewing condition, and to subsequently test whether any significant association differed in threat versus neutral conditions.

For the first aim, we hypothesized that in both conditions, compared to participants with lower SPS dimension scores, those with higher scores would demonstrate more neural synchrony in brain regions associated with self-referential information processing (DMN), emotional response (SN), and focused attention (CEN), and that significant representation similarities would be greater in the threat than in the neutral movie condition. We hypothesized that these synchrony differences would be associated with a greater activation difference between the neutral and threat conditions in participants with a higher SPS dimension score. Second, we hypothesized that in both conditions, compared to participants with lower SPS dimension scores, those with higher scores would have higher ISFC between SN and CEN, and between SN and DMN, and higher ISFC within the SN and CEN; and that significant ISFC-SPS correlations would be greater in the threat than in the neutral movie condition.

## 2. METHODS

### 2.1 Participants

Demographic, SPS, and MRI data were collected from participants as part of the ongoing Healthy Brain Study, a longitudinal population-based project acquiring data from healthy people aged 30-39 years (Healthy Brain Study consortium et al., 2021). Participants were recruited from the general population in Nijmegen, the Netherlands. All participants provided informed, written consent prior to data collection in accordance with experimental procedures approved by the Institutional Review Board of Radboud University Medical Center (reference number: 2018–4894) and the latest revision of the Declaration of Helsinki (World Medical Association, 2013). Demographics are reported in **Table 1**.

**Table 1.** Study sample demographics (N=238). SPSQ-SF: 24-item short form of the Sensory Processing Sensitivity Questionnaire (De Gucht & Woestenburg, 2024)

|  | Mean or<br>count | Standard<br>deviation or<br>percent |
| --- | --- | --- |
| Age (years) | 33.86 | 2.79 |
| Female sex (n, %) | 138 | 57.98% |
| Head motion (relative root-mean-square) | 0.12 | 0.05 |
| **<u>SPSQ-SF raw score</u>** |  |  |
| Sum | 107.53 | 18.52 |
| Positive dimension | 81.72 | 13.51 |
| Negative dimension | 25.81 | 8.36 |
| **<u>Education (n, %)</u>** |  |  |
| Pre-vocational secondary | 3 | 1.26% |
| Senior general secondary / pre-university secondary | 7 | 2.94% |
| Secondary vocational | 33 | 13.87% |
| Higher professional | 104 | 43.70% |
| University | 76 | 31.93% |
| Other | 4 | 1.68% |
| Did not receive any | 1 | 0.42% |
| Would not state | 10 | 4.20% |

Participants were required to be aged 30-39 and reside in the Nijmegen region, Netherlands. Exclusion criteria included: below Dutch B1 proficiency, prior history of significant psychiatric or neurological illness, neurological disease, current neuropharmaceutical medication, contraindication for MRI, previous brain surgery, and claustrophobia. Participants were assessed at three times over the course of one year: assessment 1 (A1), four months later (assessment 2; A2), and another four months later (assessment 3; A3) (**Figure 1**). A full description of the study design is available in (Healthy Brain Study consortium et al., 2021).

**Figure 1.**
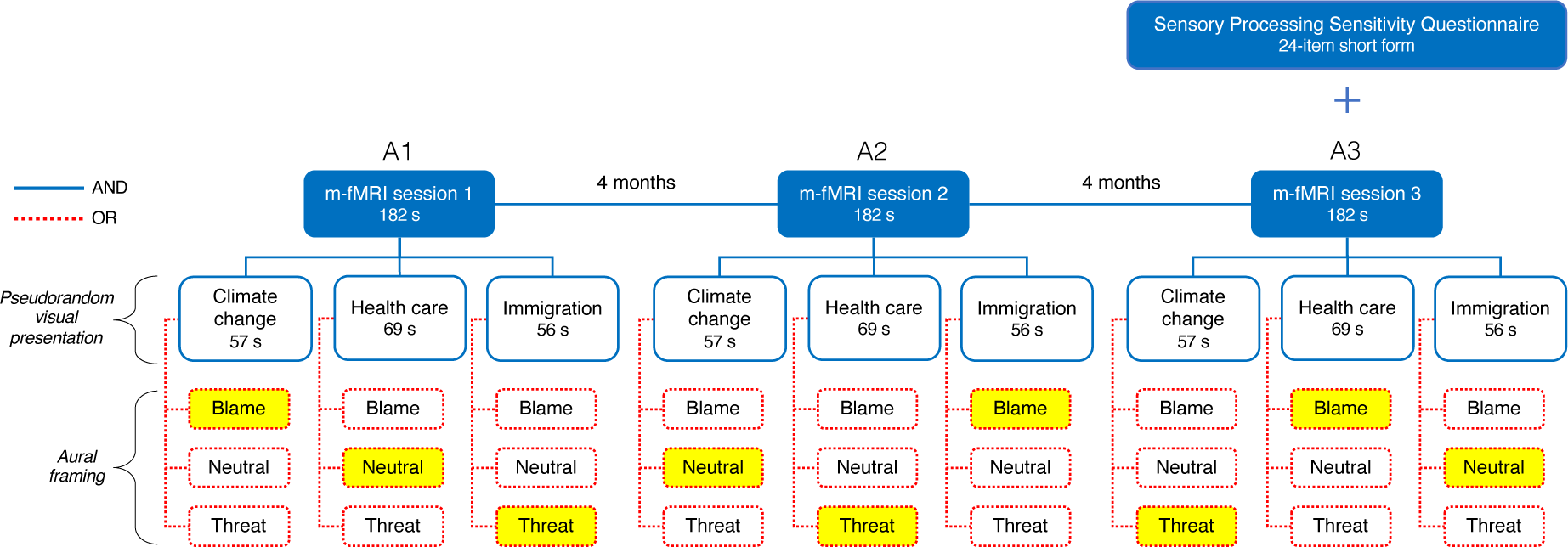
Data acquisition and task design from the Healthy Brain Study (Healthy Brain Study consortium et al., 2021). The sequence of yellow-colored boxes from left to right is an example of what one participant saw.

### 2.2 Questionnaire

At A3, participants completed the 24-item short form of the Sensory Processing Sensitivity Questionnaire (SPSQ-SF). These questionnaire items originated from (De Gucht & Woestenburg, 2024) which included 26 items. However, this short form was still under development at the time the Healthy Brain Study began, so only 24 of the 26 items were included. The 24-item version has been shown to have good psychometric properties and validated in an independent sample which overlaps with ours here (Damatac et al., 2023).

Our analyses here include SPSQ-SF data from 238 participants. To preserve individual differences in SPS, we used raw item-wise scores. In exploratory analyses, the positive and negative dimensions showed correlations in opposite directions, prompting us to focus on these two dimensions rather than subscale or total scores. We calculated the sum scores for positive (Cronbach’s α=0.52) and for negative (Cronbach’s α=0.75) dimensions (**Figure S1**).

### 2.3 MRI acquisition

Data were acquired at the Donders Institute for Brain, Cognition and Behaviour (Nijmegen, The Netherlands) with two nearly identical 3 Tesla MRI scanners: Siemens Prisma and Siemens Prismafit. Prior to data collection, participants were randomly assigned to one of these scanners and all MRI measurements across all MRI sessions were collected using the assigned equipment. Prior to A1, each participant underwent a 15-minute practice session in a mock scanner to become acquainted with the scanner environment and learn how to minimize head movements.

T1-weighted structural scans were acquired with an MPRAGE sequence (time repetition[TR]=2000ms, time echo[TE]=2.03ms, time to inversion=880ms, flip angle=8, voxel size=1mm^3^). T2*-weighted m-fMRI scans were acquired with a multiband echo-planar imaging sequence (TR=1s, TE=34ms, flip angle=60, field-of-view[x,y,z]=182×218×182mm, acquisition matrix=104×104, voxel size=2mm^3^, multiband acceleration factor=6). Data acquisition produced 235 volumetric images per subject (66 slices/volume).

Neuroimaging data were acquired with a pseudorandom movie task design (**Figure 1**). During each movie m-fMRI session (A1: assessment 1, A2: assessment 2, A3: assessment 3), each lasting for 235 seconds total, participants watched three short movies that varied by topic (immigration, climate change, and health care), each narratively framed as blame, threat, or neutral, in a 3×3 pseudorandomized design. At each assessment, a participant viewed one movie clip from each of the three topics accompanied by one of each of the three aural frames. Thus, each topic and each frame were presented only once per assessment. Throughout all assessments, participants viewed one of nine clips total, so each topic-frame combination was only ever presented once to each participant. All movies were played with Dutch audio.

### 2.4 MRI pre-processing

The dataset was preprocessed by the Healthy Brain Study. Structural data were cleaned using FSL’s “fsl_anat,” gradient distortion correction was applied, face and ears were removed to ensure anonymity, and resulting files were used as inputs for m-fMRI pre-processing. Functional data were cleaned using FSL’s FEAT toolbox (including motion correction, field maps for distortion correction, and T1 data for non-linear registration to MNI152 space), then further denoised using FSL FIX (custom training set based on a subset of 40 random participants) (Griffanti et al., 2014; Salimi-Khorshidi et al., 2014), and gradient distortion correction. A high-pass 0.01Hz temporal filter was applied to remove low-frequency scanner drifts. Data was excluded if it included more or less than 235 fMRI timepoints and head motion >4mm, resulting in the exclusion of 33 images from 26 participants.

After receiving the movie-fMRI data from the Healthy Brain Study, we identified 7 participants who showed poor registration to the MNI space and excluded those from further analysis, resulting in a final sample of 653 images from 238 subjects. We smoothed the data with a 6mm FWHM Gaussian kernel. For each subject, we used a 2mm^3^ atlas comprising 300 parcels arranged into 7 networks (Schaefer et al., 2018; Yeo et al., 2011) to estimate functional activation. We then regressed out six motion parameters, using each of the three translations and three rotations as separate regressors in a linear regression model. Then, we *z*-scored within each session to control for session-specific effects, separated the volumes by unique frame-topic combinations (excluding volumes six seconds after movie onset and including volumes six seconds after movie ending), and concatenated volumes across sessions.

### 2.5 Analyses for aim 1: Association between m-fMRI dissimilarity and SPS dimension score distance

#### 2.5.1 Brain similarity: Inter-subject correlation

To estimate network inter-subject correlation (ISC), we first calculated ISC from individual ROIs and then averaged ROI ISCs within networks. This way ROI-specific signals are still able to contribute to the ISC calculation, even though we compute the similarity to SPS on the network level. Thus, for each subject, we calculated the Pearson correlation between each ROI pair, then converted correlation to dissimilarity (=1-correlation), resulting in 300 ROI ISC matrices. We averaged ISCs across ROIs belonging to CEN, DMN, and SN, producing three network ISC matrices.

#### 2.5.2 Questionnaire similarity: Euclidean distance

Trait similarity can be represented in various ways and the choice of distance metric influences our analyses and results based on our initial assumptions (Finn et al., 2020). Therefore, as a preliminary analysis, we assessed three distinct assumptions about the structure of inter-subject SPSQ-SF score (dis)similarity: (1) nearest neighbor: a Euclidean distance model, wherein participants with similar SPS scores are most alike regardless of their absolute position on the scale; (2) centrality: a dissimilarity model so that participants with more average scores are more alike, while those at either extreme end of the scale are less alike; and (3) convergence: a similarity model in which individuals with higher SPS are more alike and those with lower SPS are less alike. For each model, we calculated similarity between each subject pair’s positive and negative dimension scores, resulting in two inter-subject matrices per model. We then Spearman correlated each of these matrices with each of the 300 ROI (whole-brain) ISC matrices. Our first model (nearest neighbor) produced the highest number of uncorrected significant associations between SPSQ-SF similarity and whole-brain ISC throughout all m-fMRI volumes (**Figure S2**). As a result, we chose nearest neighbor (i.e., Euclidean distance) as our metric of SPSQ-SF similarity for our main analyses.

#### 2.5.3 Inter-subject representation similarity

To estimate the association between ISC and SPS, we first extracted and vectorized the lower triangle of the SPS and ISC inter-subject dissimilarity matrices. Next, we used Spearman rank correlations to estimate the association between the two dissimilarity matrices, separately for the neutral and threat condition and the positive and negative SPS dimension. To calculate the significance of the association between SPS and ISC, we performed a permutation test. We permuted the SPS scores across participants before reconstructing the SPS dissimilarity matrices. This preserved the dependence structure of the data, while eliminating any association between SPS and ISC. We repeated this permutation 10,000 times and re-calculated the correlation each time to generate a null distribution of correlation coefficients. We conducted two-sided tests by calculating the *p*-value as the proportion of correlation coefficients, obtained from the null distribution, that are as extreme as or more extreme than the absolute value of the observed Spearman correlation coefficient. We then corrected all *p*-values for multiple testing with false-discovery rate (FDR) (Benjamini & Hochberg, 1995), correcting across 12 tests. If we found a significant correlation for either fMRI dissimilarity in the neutral or threat condition in relation to SPS distance, we tested if it was statistically significantly different from that in the other condition (Steiger paired correlation comparison), and whether it was driven by any specific ROIs (FDR-corrected across the number of ROIs).

#### 2.5.4 Analyses for sub-aim 1.1: Network activation

We calculated the mean of the pairwise differences between ROI activations for a threat versus neutral condition contrasts. This analysis produced a difference value per contrast, per subject, and per ROI. Next, we averaged the differences across ROIs within each *a priori* network to generate average network contrast values. Finally, for each network, we calculated Pearson correlations across subjects: difference _contrast_ ∼ SPS _dimension_. We FDR-corrected across 6 tests.

### 2.6 Analyses for aim 2: Association between ISFC and SPS dimension score

To estimate ISFC, we used the same extracted data of ROIs grouped into three *a priori* networks from the previous analysis. Connectivity was estimated using a leave-one out procedure, where every participant’s data in each ROI was correlated with the averaged data across all other participants in all ROIs. This resulted in a ROI-ROI connectivity matrix for each participant. Next, we averaged the correlations across ROI pairs, resulting in a network-by-network connectivity matrix for all within and between network connections. We then correlated the vectorized network-by-network connectivity with positive and negative SPS dimension sum scores and FDR-corrected across 12 tests. For both between- and within-network ISFC, if we found a significant correlation for either ISFC in the neutral or threat condition in relation to SPS, we then tested whether it was significantly different from that in the other condition and whether it was driven by any specific ROI pairs (FDR-corrected across the number of ROI-pairs).

## 3. RESULTS

### 3.1 Results for aim 1: Association between m-fMRI dissimilarity and SPS dimension score distance

We investigated ISC in both threat and neutral conditions across the three networks and how it related to the positive and negative SPS dimension; in particular, whether more similar SPS scores related to more similar neural responses. Only during threat-framed movies, we found that more similar negative dimension SPS score related to more similar neural responses in the CEN (Spearman’s rank correlation coefficient[*r_s_*]=0.18, false discovery rate corrected *p*-value[*p*_fdr_]=0.041) and SN (*r_s_*=0.19, *p*_fdr_=0.039) (**Figure 2, Table S1**). There were no significant differences between the correlations in neutral versus threat conditions for both effects; condition did not modulate the significant IS-RSA effects (Steiger test statistic[*z*]=-1.63, *p*=0.104) and SN (*z*=-1.74, *p*=0.083). Lastly, we found no significant effects in relation to SPSQ-SF positive dimension score or to the DMN.

**Figure 2.**
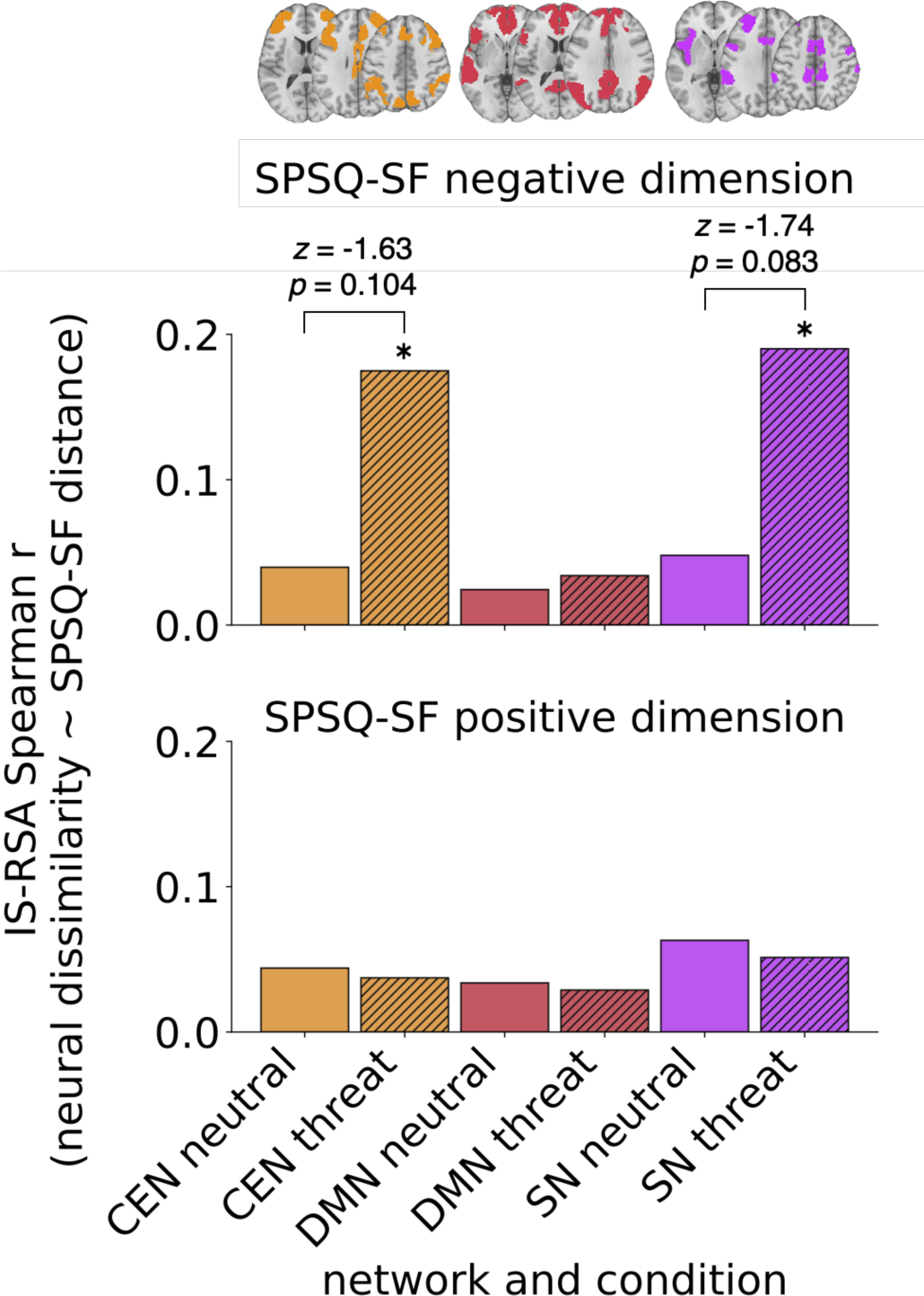
(**Top**) More inter-subject distance in the SPSQ-SF negative dimension correlated with more neural activation dissimilarity in CEN and SN and only during threat-framed movies (designated by asterisks). Network ROI anatomical locations are overlain above the bar plots. (**Bottom**) There were no significant effects in relation to SPSQ-SF positive dimension score. Bar heights: correlation values between neural ISC and SPSQ-SF dimension score Euclidean distance. Bars are colored according to network. Bars with diagonal stripes indicate threat condition, while bars without stripes indicate neutral condition. Results are tabulated in Table S1.

To follow up on this, we investigated whether these effects could be attributed to specific ROIs within these networks. We observed that there were consistent but small associations between negative dimension SPS and ISC in both CEN and SN and that there were no specific ROIs within these networks that drove the observed association at the network level (all *r_s_*<0.05, all *p*_fdr_>0.05; **Figure S3**).

#### 3.1.1 Results for sub-aim 1.1: Network activation

We investigated how activity was affected by framing condition and whether there are systematic increases or decreases in neural activity along the SPS scale (**Figure S4**).

Higher negative dimension SPS score related to a greater activation difference between threat and neutral conditions within CEN (*r_p_*=0.14, *p*_uncorrected_=0.032) (**Figure S5**, **Table S2**). In this network, more highly sensitive individuals (negative dimension) exhibited a greater activation difference between threat and neutral conditions compared to less sensitive individuals; specifically, the threat condition displayed higher activation than the neutral condition. However, this effect did not survive FDR-correction.

### 3.2 Results for aim 2: Association between ISFC and SPS dimension score

We next investigated ISFC in both threat and neutral conditions across the three networks and how it related to the positive and negative SPS dimension; in particular, whether higher SPS scores related to higher ISFC.

#### 3.2.1 Between networks

For all pairs of networks, we observed positive between-network ISFC during both threat and neutral conditions (Pearson’s correlation coefficient[*r_p_*]<0.35), which was not significantly modulated by condition across all participants (−0.21<*z*<-0.09, *p*>0.05) (**Figure 3**).

**Figure 3.**
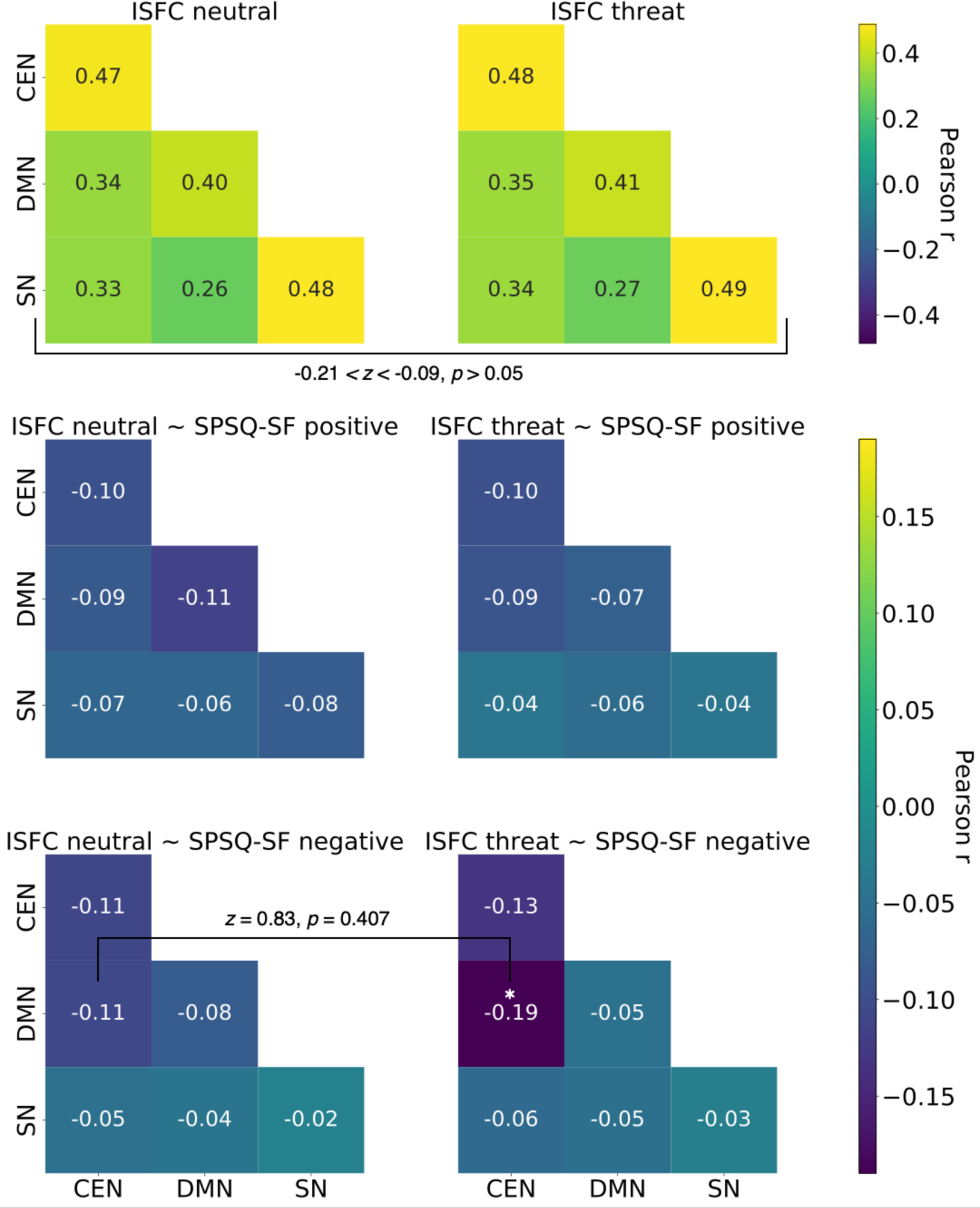
Inter-subject functional connectivity (ISFC) between and within *a priori* networks irrespective of SPS (**top**), in relation to SPSQ-SF positive dimension score (**middle**), and in relation to SPSQ-SF negative dimensions score (**bottom**). Left column: ISFCs in the neutral m-fMRI condition. Right column: ISFCs in the threat condition. Color and numerical values indicate the Pearson correlation (r). Irrespective of SPS, ISFC within and between networks was positive during both neutral and threat conditions and ISFCs were not modulated by condition. Overall, reduced ISFC correlated with higher SPS scores in both dimensions and m-fMRI conditions. Reduced ISFC between CEN and DMN during threat significantly correlated with higher SPSQ-SF negative dimension score (bottom-right panel; indicated by an asterisk). This significant effect was not modulated by condition. ISFC results are tabulated in Table S3.

All associations between ISFC and SPSQ-SF dimension score were negative, though not all were significant (i.e., reduced ISFC between networks always correlated with higher SPS; **Figure 3; Table S3**). Only during threat-framed movies, higher negative dimension SPS score significantly related to lower ISFC between CEN and DMN (*r_p_*=-0.19, *p*_fdr_=0.039) (**Figure 3**). In other words, during threat, more highly sensitive individuals (in the negative dimension) showed reduced functional connectivity between these brain networks than less sensitive individuals. This significant correlation between CEN-DMN connectivity and SPS score was not significantly modulated by condition (*z*=0.83, *p*=0.407).

When we explored this main between-network effect at the ROI level, we observed that, again, there was a small negative correlation shared across many pairs of ROIs that together appeared to drive the observed effects. Yet, one negative correlation was notable: we observed that ISFC between left lateral prefrontal cortex (CEN) and right dorsomedial prefrontal cortex (DMN) significantly correlated with SPSQ-SF negative dimension score (*r_p_*=-0.28, *p*_fdr_=0.044) (**Figure S6**). This suggests that these two regions may be the strongest drivers of the main between-network effect that we detected.

#### 3.2.2 Within networks

For all networks, we observed positive within-network ISFC during both threat and neutral conditions, which was not significantly modulated by condition across all participants (−0.214<*z*<-0.091, *p*>0.05) (Figure 4).

Only during threat-framed movies, higher negative dimension SPS score related to reduced functional connectivity within CEN (*r_p_*=-0.132, *p*_uncorrected_=0.042) (**Table S3**). During threat, more highly sensitive individuals (in the negative dimension) exhibited less functional connectivity in this network compared to less sensitive individuals. However, this effect did not survive FDR-correction. Though none were significant, all associations between ISFC and SPSQ-SF dimension score were negative (i.e., reduced ISFC within networks always correlated with higher SPS; Figure 4**; Table S3**).

## 4. DISCUSSION

We have provided a comprehensive characterization of the neural synchrony, activation, and between- and within-network functional connectivity features of SPS in a large cross-sectional sample of healthy adults. We observed that threatening or stressful environmental stimuli had a greater impact on individuals who scored higher in SPS compared to those who scored lower. Notably, we consistently found neural associations with only the negative dimension of SPS, and CEN appears to be the most consistently related to SPS during threat (i.e., while watching threat-framed movies). This study contributes to current evidence for neurobiological mechanisms related to SPS in response to environmental stimuli.

### 4.1 Differences in the CEN may play a key role in shaping environmental perception for highly sensitive individuals

Overall, our results support previous studies concluding that individuals with higher sensitivity are more affected by highly emotional or threatening stimuli, and we notably establish a connection between SPS and neural response during threat specifically in the CEN. Although we hypothesized SPS-related differences in all three *a priori* networks, this was the only network that had consistent SPS-related effects in the form of altered neural ISC (higher CEN synchrony related to higher SPS score), activation (higher threat activation related to higher SPS score), and within and between-network ISFC (reduced functional connectivity within CEN, and between CEN and DMN related to higher SPS score). Although not all these effects survived multiple comparison correction, the consistent effects in the CEN in all analyses do suggest that it plays a central role. Overall, in response to threat, our findings indicate that either the CEN is most strongly modulated by SPS, or this network exhibits a stronger response to threat in individuals with higher SPS along the negative dimension.

The CEN involves frontal and parietal regions responsible for cognitive control, emotion regulation, and working memory manipulation (Niendam et al., 2012), suggesting potential SPS-related differences in resource allocation for processing threatening stimuli. Initially, higher activation but reduced connectivity in the CEN with higher SPS may seem counterintuitive; however, it has been shown that decreases in connectivity often accompany increased activation (Cole et al., 2021). Previous research has shown that positive childhood experiences and current exposure to negative stimuli in highly sensitive women increased fMRI activation in CEN brain areas (Acevedo et al., 2017), suggesting that SPS-related effects in this network are influenced by both past experiences and current negative environmental stimuli. This is in line with our findings that the effects of SPS are only observed in the threat condition. Stress research demonstrated a shift in neural resource allocation during acute stress response, characterized by heightened connectivity within SN accompanied by reduced connectivity within CEN, an adaptive redistribution that enables flexible responses to fluctuating environmental stimuli (Hermans et al., 2011, 2014). Taken together with our results, this suggests that, for highly sensitive people, a heightened neural responsiveness in the CEN may contribute to increased vigilance and arousal in response to perceived threatening stimuli, potentially leading to heightened emotional reactivity and susceptibility to stress. Additionally, adaptive neural resource redistribution may have varied effects on individuals with high SPS, depending on environmental stimuli. Some may benefit from enhanced adaptability and responsiveness, while others may experience maladaptive responses such as hyper-reactivity or cognitive overload. We tentatively propose that the CEN may involve regions where differential resource reallocation occurs during encounters with threatening or stressful stimuli in highly sensitive individuals.

We selected *a priori* networks based on Homberg & Jagiellowicz (2021) neural model, which proposed increased within-network connectivity in the CEN when highly sensitive individuals process threat-related cues. However, contrary to this hypothesis, we observed decreased connectivity but increased activation within CEN during threat in more highly sensitive people. This suggests that individual regions within CEN may show heightened responsiveness to threat (activation) while simultaneously different parts of the CEN are less coordinated in their activity (connectivity). It is difficult to determine how this would relate to the ability of high SPS individuals to respond to threat effectively. On one hand, reduced coordination within CEN could reflect a more selective neural response or be related to increased integration of information between CEN and other functional networks not included in this study. Alternatively, the altered connectivity response may be related to reduced gray matter volume observed in the right dorsal anterior cingulate cortex of the CEN in individuals with higher SPS (Wu et al., 2021), indicating potential network integration or efficiency differences, perhaps directly caused by gray matter changes, or serving as a compensatory mechanism.

#### 4.1.2 Highly sensitive individuals may process threat differently

Our study additionally sheds light on how highly sensitive individuals may process threat differently. The Homberg & Jagiellowicz (2021) model suggests that highly sensitive individuals exhibit increased connectivity between the CEN and SN, as well as between SN and DMN, when processing threat-related cues. Contrary to these hypotheses, our results indicate that communication between these networks may not directly correlate with SPS in the observed experimental conditions. Instead, we observed SPS-related effects within the CEN and its connectivity with the DMN, which were reduced in participants with higher SPS specifically during threat. Our results did show partial support for the hypothesis that effects would be specifically observed during threat-framed movies, rather than neutral-framed ones. Additionally, comparing threat to neutral conditions, all connectivity values (except within DMN) decreased, suggesting a heightened threat response associated with higher SPS that trends towards further reduced connectivity. This implies a distinct neural response to threat stimuli in more highly sensitive individuals. However, our interpretations remain speculative as we did not observe significant modulation of effects by aural framing condition. Nevertheless, the consistency of SPS-related neural alterations exclusively during threat conditions suggests differences in processing threat, possibly reflecting adaptive mechanisms or heightened sensitivity to environmental cues.

In individuals with higher SPS, reduced connectivity within the CEN and its links with the DMN, particularly evident during threat processing, suggest a more selective neural response to threatening cues, possibly reflecting adaptive mechanisms (Homberg & Jagiellowicz, 2021). This reorganization of neural networks may prioritize emotional processing and introspective thought, enhancing their ability to navigate emotional challenges (Acevedo et al., 2014). It has previously been suggested that altered DMN activation and reduced DMN-CEN connectivity may complementarily facilitate emotional processing and self-referential thought, potentially aiding in emotional regulation (Buhle et al., 2014; Cocchi et al., 2013). Dysregulation in the interplay between these networks during threat may contribute to difficulties in attentional shifting, emotion regulation, or overgeneralization of threat (Lee & Siegle, 2012). The increased synchrony within the SN that we observed in highly sensitive individuals may reflect enhanced threat detection and processing, aligning with the notion of heightened sensitivity as an adaptive trait (Acevedo et al., 2021b; Pluess, 2015).

### 4.2 Each SPSQ-SF dimension may relate to distinct neurological mechanisms

Importantly, we linked the negative dimension of SPS to brain responses to threat, indicating that individuals with higher negative dimension SPS may process such stimuli differently, potentially involving altered cognitive resource allocation, particularly within the CEN. While we expected neural alterations in both SPS dimensions, we only observed associations with the negative dimension, supporting existing literature that distinguishes between the two dimensions (Damatac et al., 2023; De Gucht et al., 2022; De Gucht & Woestenburg, 2024) and emphasizing the need to separate them when studying environmental responses. In our study, neural response to threat-related stimuli was mainly linked to the negative aspects of SPS (subscales *emotional and physiological reactivity* and *sensory discomfort*), reflecting distinct processes underlying positive and negative high sensitivity. Psychologically, individuals scoring high on the negative dimension of SPS may display traits such as more emotional reactivity, stress sensitivity, and aversive reactions to sensory stimuli, which are traits typically associated with negative affect, increased stress levels, and a tendency to feel overwhelmed by environmental stimuli (De Gucht et al., 2022). These traits could physiologically manifest through unique patterns of arousal and reactivity within the autonomic nervous system. Specifically, individuals with higher scores on the negative dimension may demonstrate greater sympathetic arousal, resulting in amplified physiological reactions to stressors and aversive stimuli (Fechir et al., 2010). These physiological differences may then be reflected in the altered neural responses observed in fMRI blood oxygen level-dependent signals.

### 4.3 Strengths and limitations

Our study’s focus solely on threat-related stimuli limits generalizability compared to a broader range of emotional experiences; absence of positive stimuli might account for lack of effects associated with positive SPS dimension. The repeated visual presentation of movies across varied aural framing conditions, albeit pseudorandomized, could have introduced familiarity and exposure biases, potentially impacting neural responses regardless of aural conditions. Furthermore, our conclusions are constrained by an *a priori* search space; a whole brain analysis could offer a more comprehensive examination of SPS-related neural activation, though there are arguments favoring a hypothesis-based approach. Nonetheless, this study provides valuable insights into SPS neural underpinnings, laying a foundation for further exploration of individual brain differences related to SPS.

Our study’s use of IS-RSA, activation, and ISFC measures enhanced understanding of how SPS relates to neural processing, with added specificity from incorporating SPS dimensionality, especially the negative dimension. The larger, population-based sample improved generalizability compared to prior studies with smaller or specific samples. Focusing on healthy adults aged 30-39 captured a stage after major developmental brain changes and before aging or neurodegenerative disease onset, increasing relevance while excluding neurodevelopmental effects. Finally, m-fMRI improved ecological validity by simulating more realistic situations, providing natural context for studying neural processing.

## 5. CONCLUSION

In conclusion, inter-subject representation similarity, activation, and inter-subject functional connectivity analyses in a large sample demonstrated distinct SPS dimension associations with specific brain networks in different emotional contexts. We uncovered relationships between the negative dimension of SPS and neural synchrony, activation, and connectivity, suggesting that highly emotional or threatening stimuli exert a greater impact on individuals with high SPS. This comprehensive characterization of neural responses points to a neurobiological basis for SPS, emphasizing the distinct neural responses to threat or stress in individuals with a higher negative dimension of SPS. The observed neural patterns shed light on the intricate interplay between the CEN, DMN, and SN, offering valuable insights into the neural substrates of SPS in response to environmental features.

## Abbreviations

(A1): first assessment
(A2): second assessment
(A3): third assessment
(CEN): central executive network
(DMN): default mode network
(ISC): inter-subject correlation
(ISFC): inter-subject functional connectivity
(IS-RSA): inter-subject representation similarity analysis
(m-fMRI): movie functional magnetic resonance imaging
(ROI): region-of-interest
(SN): salience network
(SPS): sensory processing sensitivity
(SPSQ-SF): 24-item Sensory Processing Sensitivity Questionnaire Short Form

## Acknowledgements

The authors thank everyone who participated in this study, all the researchers who collected the data, and the Healthy Brain Study team and scientific board. We also thank Elisa van der Plas, Karin Heidlmayr, and Alan Sanfey for their guidance in how the movie fMRI data were structured.

## STATEMENTS

### Data and code availability

The original data supporting the findings of this study are available upon request to the Healthy Brain Study Consortium (Healthy Brain Study consortium et al., 2021). Due to third-party constraints, the authors cannot publicly share the data. Researchers interested in accessing the data may contact the Healthy Brain Study Consortium through their website at www.healthybrainstudy.nl. Requests will be subject to review and approval by the consortium to ensure compliance with ethical standards and data protection regulations. Approved requests will then require a formal data sharing agreement.

Prior to accessing data, the authors pre-registered their study on the Open Science Framework (<u>osf.io/w4yax</u>). All pre-processing and analysis scripts are available on GitHub (<u>bit.ly/sps_mfMRI</u>). Data analysis was performed with Python version 3.6.2 (Van Rossum & Drake, 2009).

### Ethics approval

The authors assert that all procedures contributing to this work comply with the ethical standards of the relevant national and institutional committees on human experimentation and with the Helsinki Declaration of 1975 as revised in 2008 (World Medical Association, 2013).

### Open researcher and contributor ID (ORCID)

C. G. Damatac: 0000-0001-9495-5243
J. R. Homberg: 0000-0002-7621-1010
T. E. Galesloot: 0000-0002-2717-3679
L. Geerligs: 0000-0002-1624-8380
C. U. Greven: 0000-0003-2402-3074

### Contributor roles taxonomy (CRediT)

Conceptualization: LG, CG, CD Data curation: CD

Formal analysis: CD

Funding acquisition: LG, CG, JH, TG

Investigation: CD

Methodology: LG, CG, CD

Project administration: CD

Resources:

Software:

Supervision: LG, CG

Validation:

Visualization: CD

Writing – Original Draft: CG, CD

Writing – Review & Editing: LG, CG, JH, CD

---

**Figure S1.**
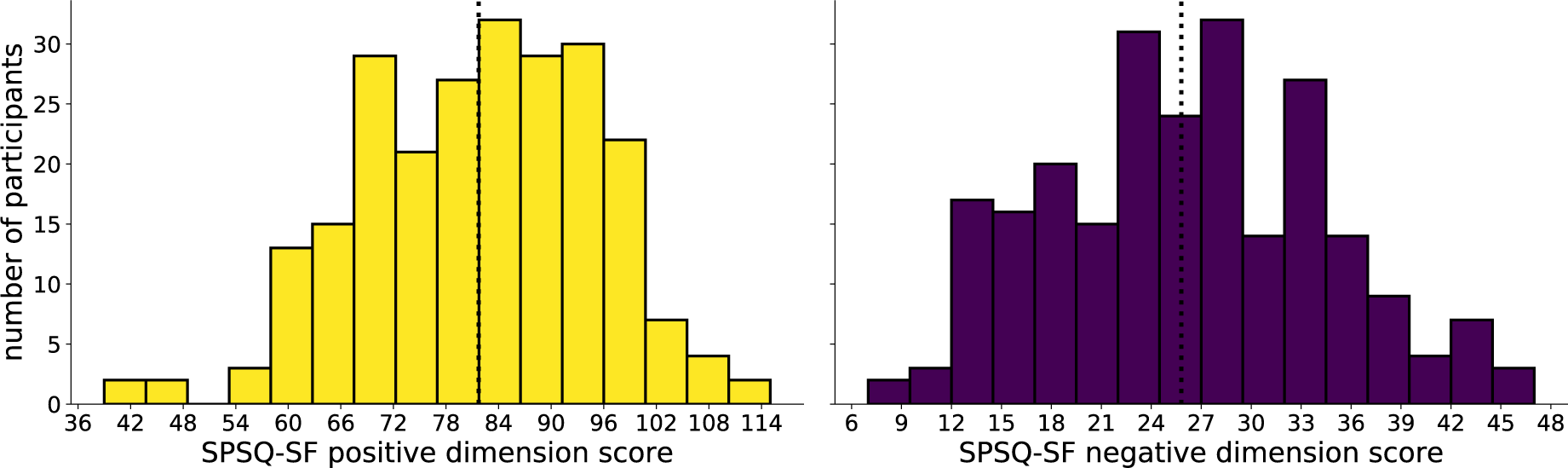
SPSQ-SF score histograms for positive (left) and negative (right) dimensions (x-axis) by the numerical count of participants (y-axis). Black vertical dotted lines represent the mean scores for either dimension.

**Figure S2.**
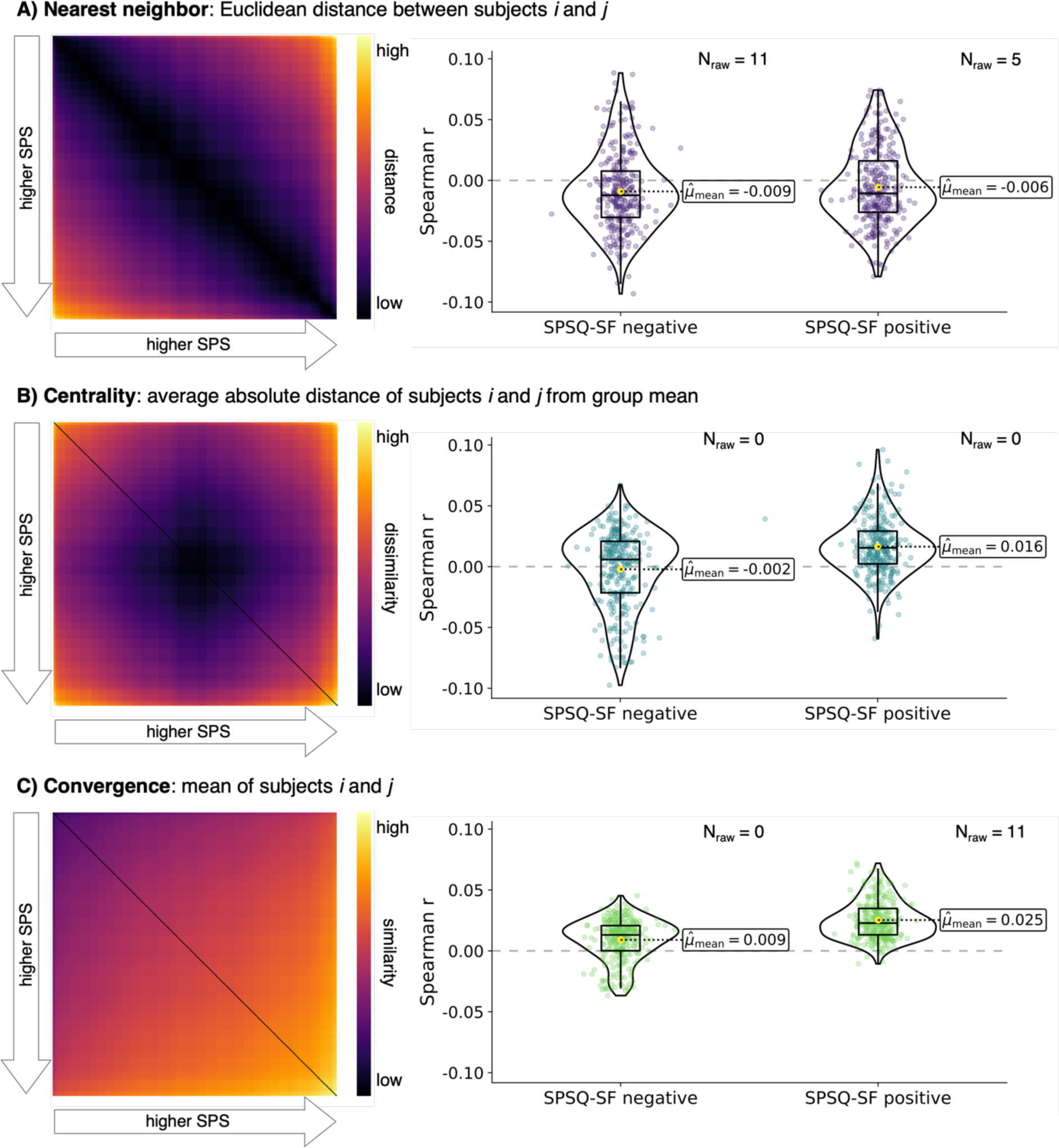
Similarity models of SPSQ-SF that we tested against whole-brain ISC for all m-fMRI volumes combined. **(A)** Euclidean distance: √((x*_i_* – x*_j_*)^2^ + (y*_i_* – y*_j_*)^2^); **(B)** Centrality: (|x*_i_* – μ| + |x*_j_* – μ|) / 2; **(C)** Convergence: mean(*i*, *j*). The box plots overlain on violin plots on the right show the Spearman correlation (*r*) between ISC and SPSQ-SF similarity for all 300 ROIs. Each point represents one ROI. N_raw_: number of uncorrected significant correlations (*p*_uncorrected_<0.05). The yellow dots represent the mean Spearman *r* across ROIs.

**Figure S3.**
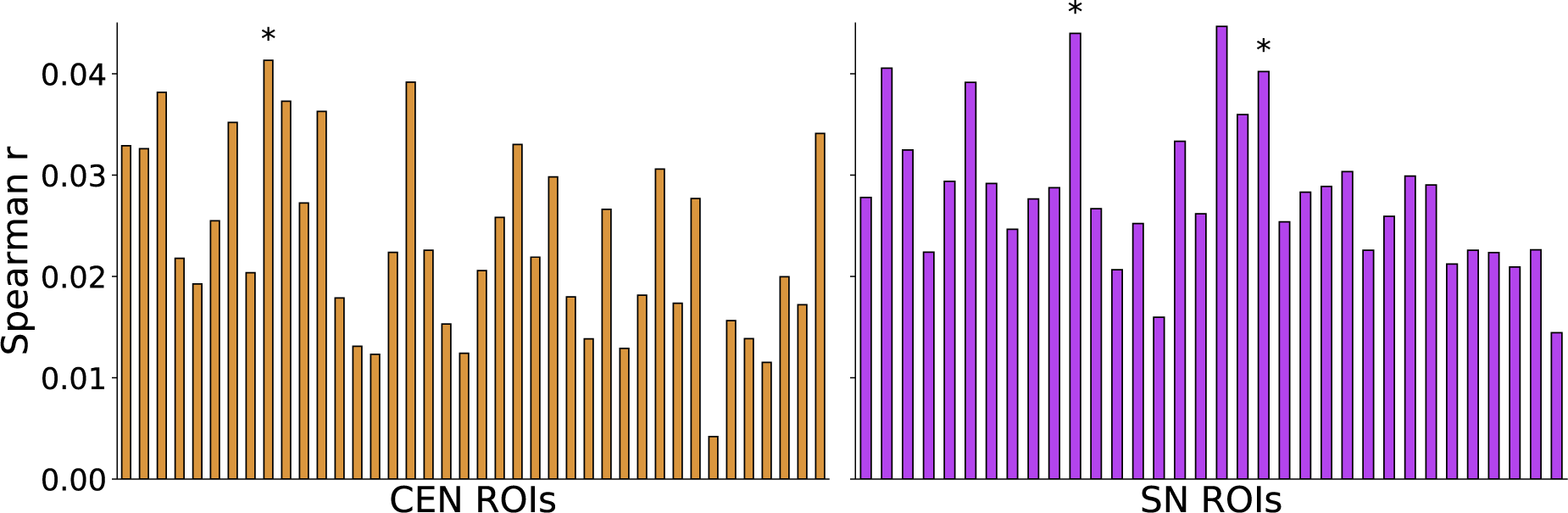
ROI-specific inter-subject representation similarity analysis results during the threat condition for **(left)** CEN and **(right)** SN. We observed positive and consistent but small associations between negative dimension SPS and ISC in both networks. Asterisks denote a significant uncorrected correlation for that ROI, but none survived multiple testing correction across the number of ROIs in each network. While there were individual ROIs showing significant correlations between SPS and ISC, these correlations were small and not consistently observed across all ROIs within the networks. Therefore, it suggests that the observed association between negative dimension SPS and ISC is more of a general pattern across the networks rather than being driven by specific ROIs.

**Figure S4.**
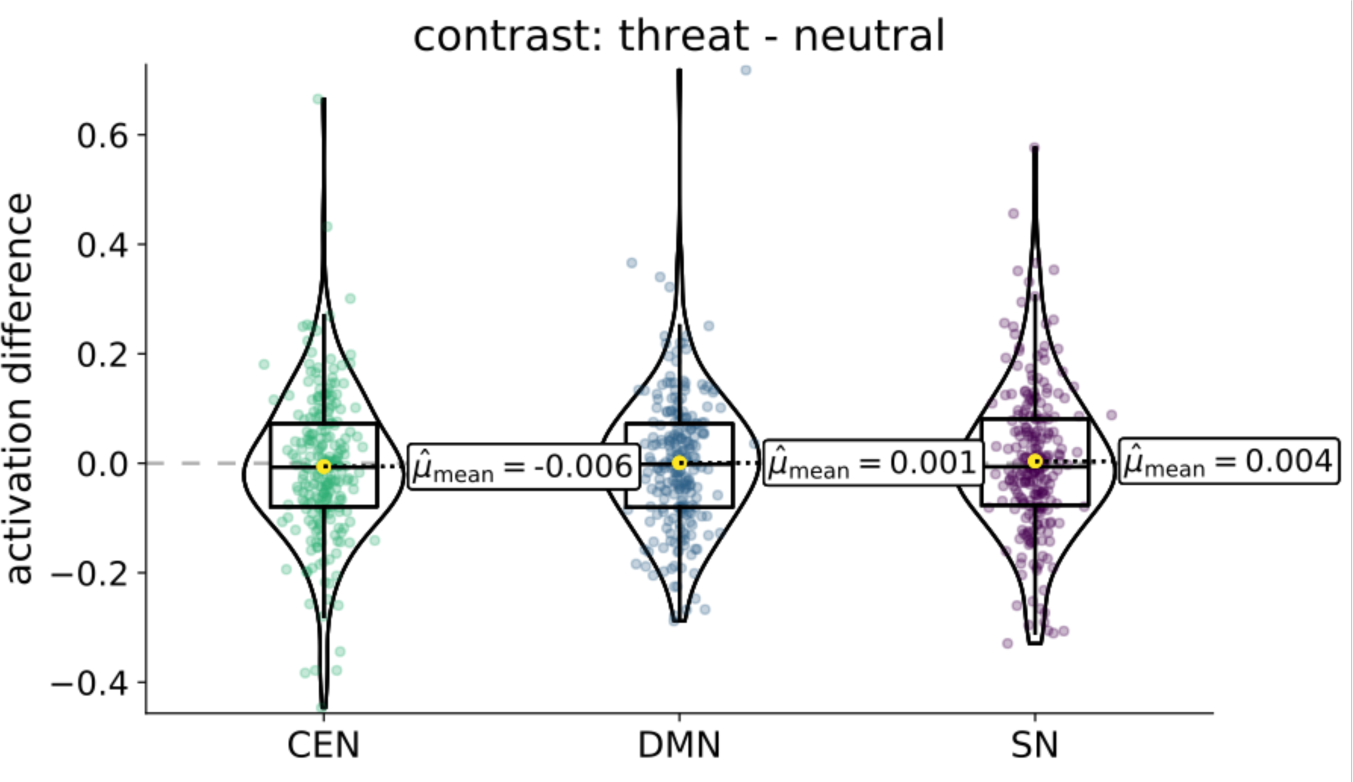
Network activation contrast differences for threat – neutral (irrespective of SPS. Network activation difference was first calculated at the ROI level and then the activations (or differences) were averaged across within-network ROIs. Box plots are overlain on violin and jittered scatterplots, wherein each point represents a participant, while labeled yellow dots indicate averages across participants.

**Figure S5.**
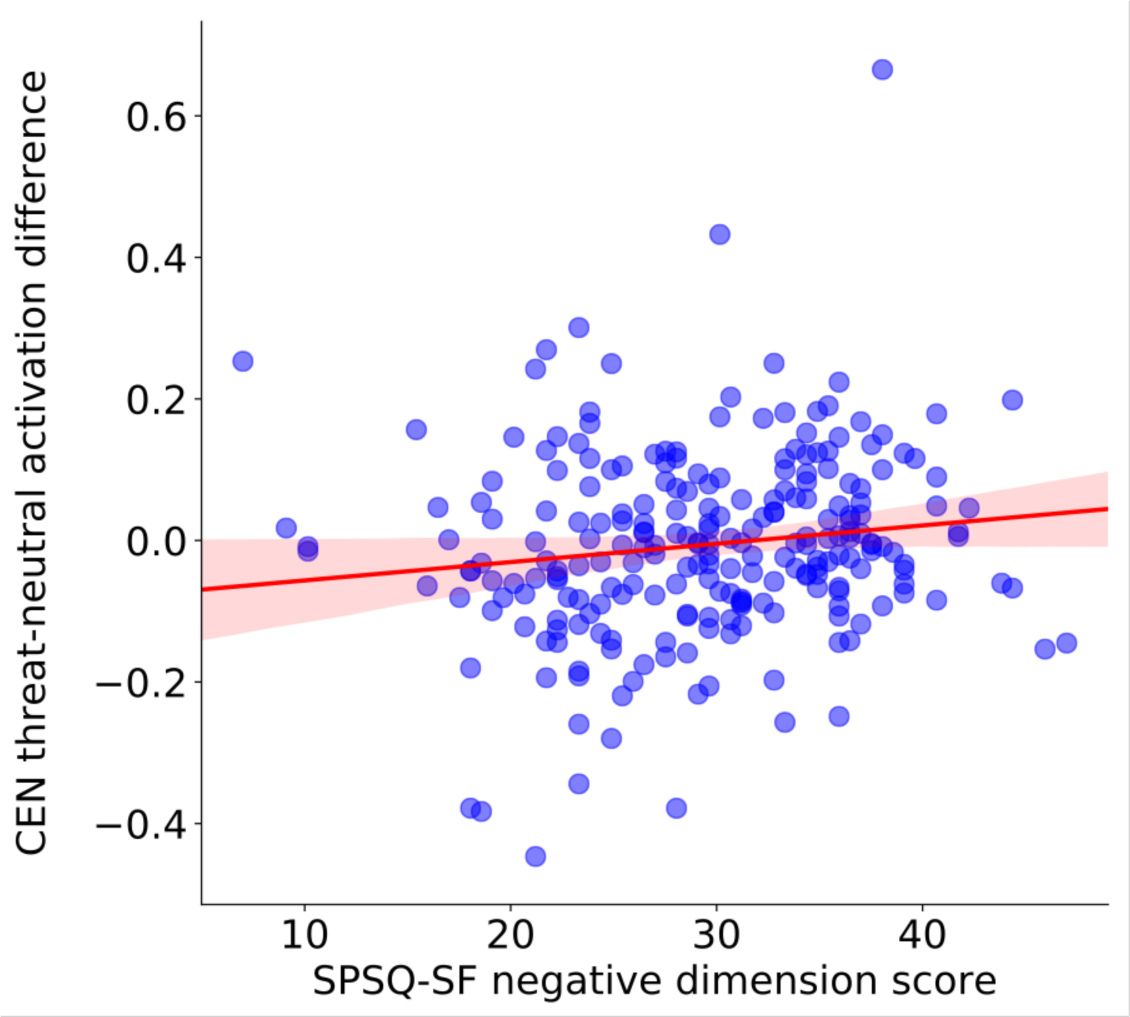
Correlation between SPSQ negative dimension score and threat-neutral activation difference in the CEN. In people with higher negative dimension SPS, CEN had higher activation in the threat condition compared to neutral. This effect did not survive false-discovery rate correction. Tabulated results are in Table S2. Each point represents one participant (darker areas indicate overlaps), and the red line represents the Pearson correlation with a 95% confidence interval.

**Figure S6.**
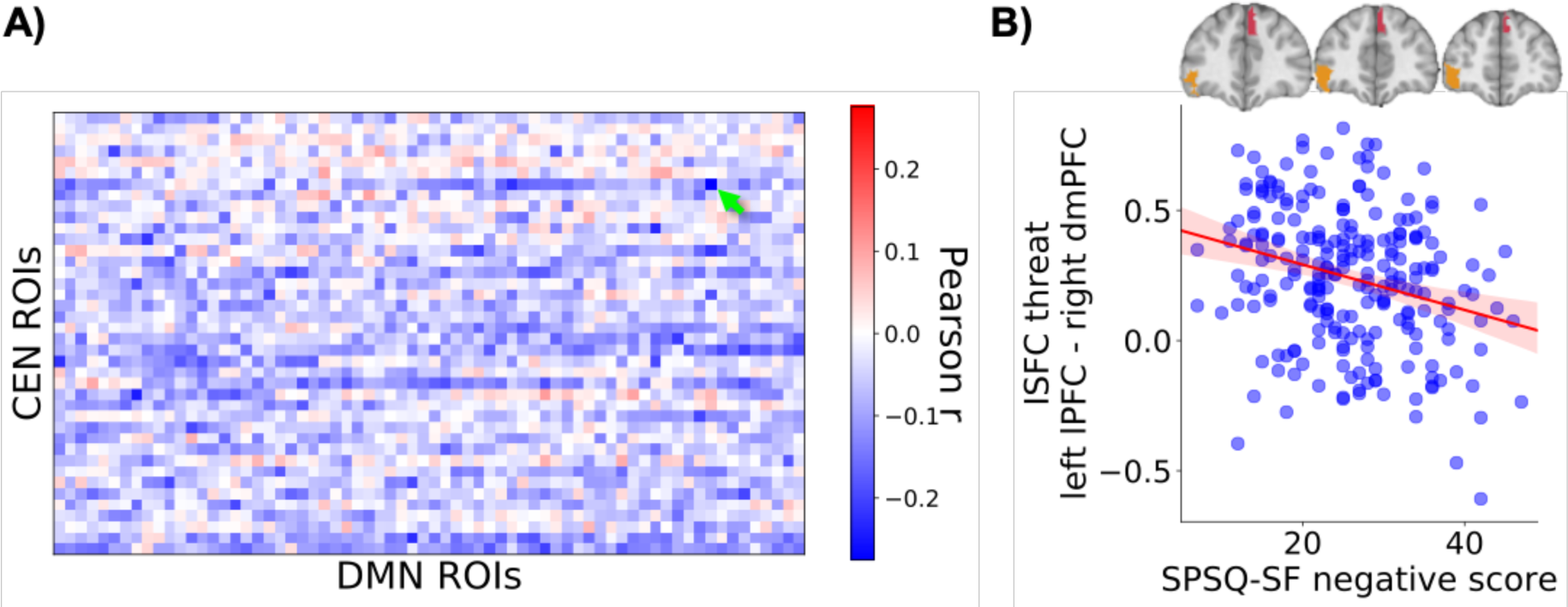
**(A)** ROI-specific CEN-DMN network inter-subject functional connectivity (ISFC) for the threat condition in association with SPSQ-SF negative dimension score. Green arrow: After false-discovery rate (fdr) correction across all CEN ROI-DMN ROI connectivities, the threat condition ISFC between left lateral prefrontal cortex (lPFC) and right dorsomedial prefrontal cortex (dmPFC) in association with SPSQ-SF negative dimension score remained significant (*r_p_*=-0.275, *p*_fdr_=0.044). **(B)** In the threat condition, higher SPS in the negative dimension related to reduced connectivity between these two ROIs. Each point represents one participant (darker areas indicate overlaps), and the red line depicts a Pearson correlation line with a 95% confidence interval. ROI anatomical locations are shown above the scatterplot (orange: left lPFC; pink: right dmPFC).

**Table S1.** Network inter-subject representation similarity analysis results. Tabulated values for Figure 3. Significant associations between m-fMRI aural framing condition functional network dissimilarity and SPSQ-SF dimension score Euclidean distance are marked with an asterisk.

| Network | Condition | SPSQ dimension | Spearman $r$ | $p$ permuted | $p$ permuted FDR-corrected |
| --- | --- | --- | --- | --- | --- |
| CEN | neutral | negative | 0.04 | 0.541 | 0.708 |
| CEN | neutral | positive | 0.04 | 0.499 | 0.708 |
| CEN | threat | negative | 0.18 | 0.007 * | 0.041 * |
| CEN | threat | positive | 0.04 | 0.567 | 0.708 |
| DMN | neutral | negative | 0.02 | 0.708 | 0.708 |
| DMN | neutral | positive | 0.03 | 0.604 | 0.708 |
| DMN | threat | negative | 0.03 | 0.601 | 0.708 |
| DMN | threat | positive | 0.03 | 0.658 | 0.708 |
| SN | neutral | negative | 0.05 | 0.461 | 0.708 |
| SN | neutral | positive | 0.06 | 0.333 | 0.708 |
| SN | threat | negative | 0.19 | 0.003 * | 0.039 * |
| SN | threat | positive | 0.05 | 0.431 | 0.708 |

**Table S2.** Correlation between difference_threat-neutral_ and SPSQ-SF dimension score across subjects. In people with higher negative dimension SPS, CEN had higher activation in the threat condition compared to neutral. This effect did not survive false-discovery rate (fdr) correction.

| Network | SPSQ dimension | Pearson $r$ | $p$ uncorrected | $p$ fdr-corrected |
| --- | --- | --- | --- | --- |
| CEN | negative | 0.14 | 0.032 * | 0.191 |
|  | positive | 0.004 | 0.955 | 0.955 |
| DMN | negative | -0.05 | 0.429 | 0.617 |
|  | positive | 0.04 | 0.514 | 0.617 |
| SN | negative | 0.06 | 0.368 | 0.617 |
|  | positive | 0.06 | 0.347 | 0.617 |

**Table S3.** Between- and within-network inter-subject functional connectivity (ISFC) associations with SPSQ-SF dimension score for each m-fMRI aural framing condition. Significant associations between m-fMRI aural framing condition ISFC and SPSQ-SF dimension score are marked with an asterisk.

| | Condition | ISFC | SPSQ dimension | Pearson $r$ | $p$ uncorrected | $p$ fdr-corrected |
| --- | --- | --- | --- | --- | --- | --- |
| **Between-network ISFC** | neutral | CEN-DMN | negative | -0.11 | 0.096 | 0.509 |
|  | neutral | CEN-DMN | positive | -0.09 | 0.187 | 0.509 |
|  | neutral | CEN-SN | negative | -0.05 | 0.443 | 0.509 |
|  | neutral | CEN-SN | positive | -0.07 | 0.312 | 0.509 |
|  | neutral | DMN-SN | negative | -0.04 | 0.506 | 0.509 |
|  | neutral | DMN-SN | positive | -0.06 | 0.326 | 0.509 |
|  | threat | CEN-DMN | negative | -0.19 | 0.003 * | 0.039 * |
|  | threat | CEN-DMN | positive | -0.09 | 0.168 | 0.509 |
|  | threat | CEN-SN | negative | -0.06 | 0.357 | 0.509 |
|  | threat | CEN-SN | positive | -0.04 | 0.509 | 0.509 |
|  | threat | DMN-SN | negative | -0.05 | 0.426 | 0.509 |
|  | threat | DMN-SN | positive | -0.06 | 0.375 | 0.509 |
| **Within-network ISFC** | neutral | CEN | positive | -0.10 | 0.143 | 0.342 |
|  | neutral | CEN | negative | -0.11 | 0.092 | 0.342 |
|  | neutral | DMN | positive | -0.11 | 0.077 | 0.342 |
|  | neutral | DMN | negative | -0.08 | 0.212 | 0.377 |
|  | neutral | SN | positive | -0.08 | 0.220 | 0.377 |
|  | neutral | SN | negative | -0.02 | 0.702 | 0.702 |
|  | threat | CEN | positive | -0.10 | 0.135 | 0.342 |
|  | threat | CEN | negative | -0.13 | 0.042 * | 0.342 |
|  | threat | DMN | positive | -0.07 | 0.253 | 0.380 |
|  | threat | DMN | negative | -0.05 | 0.464 | 0.614 |
|  | threat | SN | positive | -0.04 | 0.512 | 0.614 |
|  | threat | SN | negative | -0.03 | 0.670 | 0.702 |

